# Structural basis for variable IgE reactivities of Cor a 1 hazelnut allergens

**DOI:** 10.1101/2020.11.27.400978

**Authors:** Sebastian Führer, Anna S. Kamenik, Ricarda Zeindl, Bettina Nothegger, Florian Hofer, Norbert Reider, Klaus R. Liedl, Martin Tollinger

**Affiliations:** Institute of Organic Chemistry and Center for Molecular Biosciences Innsbruck (CMBI), University of Innsbruck, Innrain 80/82, 6020 Innsbruck, Austria; Institute of General, Inorganic and Theoretical Chemistry and Center for Molecular Biosciences Innsbruck (CMBI), University of Innsbruck, Innrain 80/82, 6020 Innsbruck, Austria; Department of Dermatology, Venerology and Allergology, Medical University of Innsbruck, Anichstraße 35, 6020 Innsbruck, Austria

## Abstract

A major proportion of allergic reactions to hazelnuts (*Corylus avellana*) are caused by immunologic cross-reactivity of IgE antibodies to pathogenesis-related class 10 (PR-10) proteins. Intriguingly, the four known isoforms of the hazelnut PR-10 allergen Cor a 1, denoted as Cor a 1.0401-Cor a 1.0404, share sequence identities exceeding 97 % but possess different immunologic properties. In this work we describe the NMR solution structures of these proteins and provide an in-depth study of their biophysical properties. Despite sharing highly similar three-dimensional structures, the four isoforms exhibit remarkable differences regarding structural flexibility, hydrogen bonding and thermal stability. Our experimental data reveal an inverse correlation between structural flexibility and IgE-binding, with the most flexible isoform having the lowest IgE-binding potential, while the isoform with the most rigid backbone scaffold displays the highest immunologic reactivity. These results point towards a significant entropic contribution to the process of antibody binding.

## Introduction

Nut allergies in Europe are predominantly related to hazelnuts (Datema et al., 2018; Datema et al., 2015) and walnuts (Lyons et al., 2020). Immunologic reactions to these food sources are triggered by several specific proteins, with a significant proportion of individuals being affected by class 10 of pathogenesis-related proteins (PR-10) (Datema et al., 2018). These allergic reactions result in particular from an initial sensitization to the major birch (*Betula verrucosa*) pollen allergen Bet v 1, a PR-10 protein. Thereafter, immunologic cross-reactivity is developed due to the strong similarity between Bet v 1 and homologous nut allergens (Ebner et al., 1995; Vieths, Scheurer, & Ballmer-Weber, 2002). Indeed, up to 70 % of all birch pollen allergic individuals are affected by such birch pollen-related food allergies (BPRFA), with hazelnuts representing one of the most prevalent triggers (Eriksson, Formgren, & Svenonius, 1982; Geroldinger-Simic et al., 2013; Geroldinger-Simic et al., 2011; K. S. Hansen et al., 2009). In affected patients, consumption of hazelnuts provokes a variety of clinical symptoms, including itching, scratching, and swelling of the mouth and throat (Mari, Ballmer-Weber, & Vieths, 2005) as well as in rare cases severe anaphylactic shocks (Kleine-Tebbe, Vogel, Crowell, Haustein, & Vieths, 2002; Le, van Hoffen, Lebens, Bruijnzeel-Koomen, & Knulst, 2013). Contrary to other food sources, industrial processing at high temperatures does not prevent IgE reactivity in hazelnut products (Wigotzki, Steinhart, & Paschke, 2001). Additionally, hazelnut allergens only decompose at much higher temperatures (Müller et al., 2000; Verhoeckx et al., 2015) than, for example, apple allergens (Somkuti, Houska, & Smeller, 2011).

PR-10 proteins, which are expressed in plants upon environmental or pathogenic stimuli, have a molecular weight of ca. 17.5 kDa and comprise about 160 amino acid residues. These proteins exhibit a canonical fold consisting of a seven stranded antiparallel β-sheet (β1 - β7) and three α-helices (α1, α2, α3). The two short, consecutive helices α1 and α2 interrupt the β-sheet between strands β1 and β2 while the long C-terminal helix α3 is located above the β-sheet, creating a large and fairly hydrophobic cavity in the protein interior (Fernandes, Michalska, Sikorski, & Jaskolski, 2013). The PR-10 proteins of the common hazel (*Corylus avellana*) can be grouped into the isoallergens Cor a 1.01 (hazel pollen) (Breiteneder et al., 1993), Cor a 1.02 and Cor a 1.03 (hazel leaf) (Hoffmann-Sommergruber et al., 1997), and Cor a 1.04 (hazelnut) (Hirschwehr et al., 1992). Interestingly, hazelnut Cor a 1.04 allergens are closely related to the birch pollen allergen Bet v 1, with amino acid sequence identities of about 83 % (Costa, Mafra, Carrapatoso, & Oliveira, 2016; Vieths et al., 2002), while hazelnut and hazel pollen allergens share sequence identities of only about 63 %. Likewise, the IgE-epitopes of the hazelnut allergens appear to show higher similarity to those of birch pollen than hazel pollen (Lüttkopf et al., 2002), and the C-terminal α-helix of hazelnut Cor a 1.04 allergens contain a main T cell epitope that is not present in birch or hazel pollen allergens (Bohle et al., 2005).

Among each other, the four isoforms of the hazelnut (Cor a 1.0401–Cor a 1.0404) share sequence identities of at least 97 %. Nevertheless, these four allergens display strikingly different IgE-binding properties in enzyme allergosorbent tests, with a proposed immunologic ranking Cor a 1.0401 > 02 > 03 > 04 (Lüttkopf et al., 2002). The structural basis for the different IgE-binding properties has remained elusive so far. In this work we present the nuclear magnetic resonance (NMR) solution structures of all four Cor a 1.04 isoforms along with in-depth experimental data regarding the structural flexibility, thermal and temporal stability and IgE-binding properties of these proteins. Our data reveal a clear inverse correlation between structural flexibility and IgE-binding, with the most flexible isoform showing the lowest potential to bind specific IgE.

## Results and Discussion

### The four Cor a 1.04 isoforms have similar structures

To investigate the four hazelnut allergens Cor a 1.0401, Cor a 1.0402, Cor a 1.0403 and Cor a 1.0404 in a comparative manner, we determined their NMR solution structures under identical experimental conditions (Figure 1). As expected, all four proteins consist of a seven stranded antiparallel β-sheet (β1-β7) and three α-helices (α1, α2, α3), which adapt the canonical PR-10 fold with helices α1 and α2 arranged in a V-shaped manner above the curved β-sheet, acting as support for the long C-terminal helix α3. The NMR structural ensembles of all four isoforms (PDB codes 6Y3H, 6Y3I, 6Y3K, 6Y3L) display high conformational homogeneity and well-defined secondary structure elements. Root-mean-square deviation (RMSD) values of the 20 lowest energy structures are 0.5-0.7 Å for heavy atoms in all cases and 0.4 Å for backbone atoms (Table 1). The four isoforms have very similar three-dimensional structures, with backbone RMSD values between them ranging from 2.3 Å to 3.8 Å, and our Cor a 1.0401 structure compares well to the Cor a 1.0401 structure that has been reported before, with a backbone amide RMSD of 4.4 Å (Jacob et al., 2019). An overlay of the lowest energy NMR structures of the four Cor a 1.04 isoforms shows that slight variations are present regarding the curvature of the β-sheet and the C-terminal helix α3. Very similar structural variations are also present between other PR-10 allergens, e.g. birch pollen and different food sources (Figure S1). Consistently, backbone RMSD values between the four Cor a 1.04 isoforms and the sensitizing allergen from birch pollen, Bet v 1, are all below 4.4 Å. In this regard, the hazelnut isoforms are comparable to other PR-10 food allergens (Fernandes et al., 2013).

**Table 1.** Statistics and summary of structure determination and refinement. ^a^ Numbers are given for all residues (Gly1 - Cys160). ^b^ Calculated for all residues as sum over r^-6. c^ Largest violation among the 20 lowest energy structures. ^d^ Obtained with the protein structure validation software (PSVS) suite (Bhattacharya et al., 2007). ^e^ All-atom clashscore, defined as number of overlaps (≥ 0.4 Å) per 1000 atoms.

|  | Cor a 1.0401 | Cor a 1.0402 | Cor a 1.0403 | Cor a 1.0404 |
| --- | --- | --- | --- | --- |
| PDB ID | 6Y3H | 6Y3I | 6Y3K | 6Y3L |
| Experimental restraints |  |  |  |  |
| total no. of NOE-based distance restraints <sup>a</sup> | 5027 | 4062 | 4396 | 2662 |
| intraresidue [$i = j$] | 1577 | 1232 | 1630 | 1175 |
| sequential [$\vert i-j\vert = 1$] | 1200 | 1133 | 1068 | 669 |
| medium range [$1 < \vert i-j\vert < 5$] | 900 | 771 | 701 | 327 |
| long range [$\vert i-j\vert \geq 1$] | 1350 | 926 | 997 | 491 |
| dihedral angle restraints | 267 | 260 | 257 | 255 |
| hydrogen bond restraints | 142 | 148 | 140 | 149 |
| total no. of restraints | 5436 | 4470 | 4793 | 3066 |
| total no. of restraints per residue | 34.0 | 27.9 | 30.0 | 19.2 |
| long range restraints per residue | 8.4 | 5.8 | 6.2 | 3.1 |
| Restraint violations <sup>b</sup> |  |  |  |  |
| average distance violation | 0.032 ±<br>0.024 Å | 0.015 ±<br>0.014 Å | 0.017 ±<br>0.010 Å | 0.020 ±<br>0.017 Å |
| maximal distance violation <sup>c</sup> | 0.17 Å | 0.092 Å | 0.045 Å | 0.15 Å |
| average dihedral angle violation | 0.39 ± 0.90° | 0.19 ± 0.17° | 0.46 ± 1.00° | 0.16 ± 0.15° |
| maximal dihedral angle violation <sup>c</sup> | 14.10° | 0.77° | 15.7° | 0.73° |
| RMSD values |  |  |  |  |
| backbone atoms | 0.4 Å | 0.4 Å | 0.4 Å | 0.4 Å |
| heavy atoms | 0.7 Å | 0.5 Å | 0.5 Å | 0.5 Å |
| bond lengths | 0.013 Å | 0.013 Å | 0.013 Å | 0.013 Å |
| bond angles | 2.4° | 2.0° | 2.1° | 2.1° |
| Ramachandran plot statistics <sup>d</sup> |  |  |  |  |
| most favored regions | 88.3 % | 96.2 % | 93.3 % | 96.2 % |
| allowed regions | 11.7 % | 3.0 % | 6.7 % | 3.3 % |
| disallowed regions | 0.0 % | 0.7 % | 0.0 % | 0.5 % |
| All-atom clashscore <sup>e</sup> | 3 | 0 | 0 | 0 |

**Figure 1.**
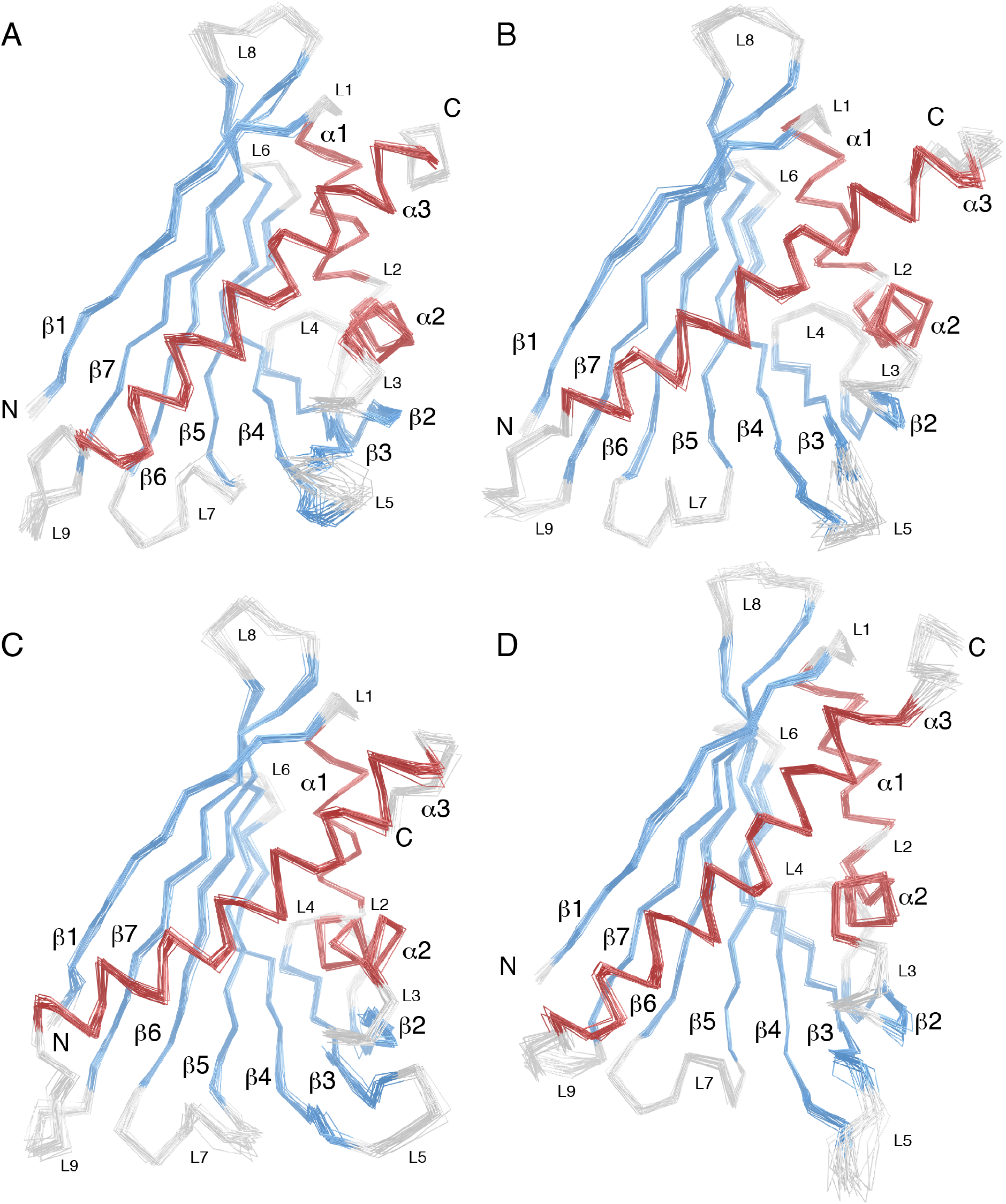
Backbone overlay of the 20 NMR solution structures with the lowest energy of the four hazelnut allergens Cor a 1.0401 (A), Cor a 1.0402 (B), Cor a 1.0403 (C), and Cor a 1.0404 (D), with the PDB codes 6Y3H, 6Y3I, 6Y3K, and 6Y3L, respectively. The secondary structure elements are defined as β1 (Val2 - Ser11), α1 (Pro15 - Leu24), α2 (Ala26 - Ala34), β2 (Thr39 - Glu45), β3 (Gly51 - Ala59), β4 (Phe64 - Asp75), β5 (Phe79 - Glu87), β6 (Glu96 - Ala106), β7 (Gly112 - Thr123), and α3 (Glu131 - His155). The α-helices are highlighted in red, β-strands in blue, and loops in light gray. Loops L1 - L9 as well as N- and C-termini are indicated.

A sequence comparison of the four Cor 1 .04 isoforms is shown in Figure 2A. Among each other, between one and six amino acid residues (at positions 4, 40, 62, 99, 130 and 158) are different, with Cor a 1.0401/Cor a 1.0404 (two residues) and Cor a 1.0402/Cor a 1.0403 (one residue) being the most similar isoform pairs. The six variable residues in the four isoforms are distributed over the protein scaffold and not in spatial proximity to each other (Figure 2B). Intriguingly, Cor a 1.0404 contains a proline residue at position 99 in the center of the β-sheet. The NMR chemical shift data clearly show that Pro99, like all other prolines, possesses trans configuration (Shen & Bax, 2010), in accordance with its location in an antiparallel β-sheet. Strand β6 (residues Glu96-Ala106) containing Pro99 and the adjacent strand β7 (Gly112-Thr123) display only slightly reduced β-strand propensities in Cor a 1.0404 when compared to the other three isoforms (Führer, Zeindl, & Tollinger, 2020).

**Figure 2.**
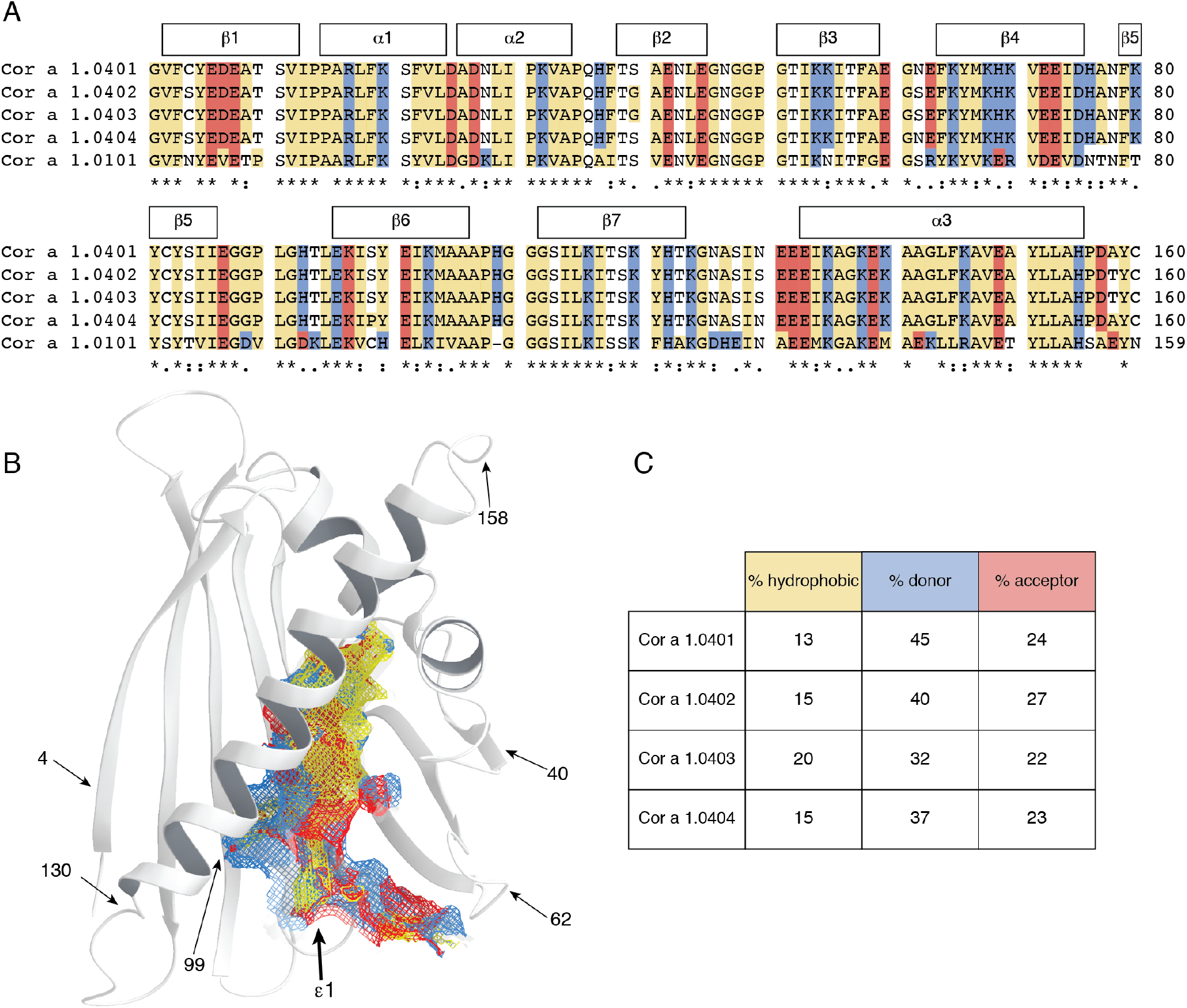
Amphiphilicity of the Cor a 1.04 internal cavity. (A) Sequence alignment of the hazelnut (Cor a 1.0401, Cor a 1.0402, Cor a 1.0403, Cor a 1.0404) and hazel pollen (Cor a 1.0101) allergens obtained with Clustal Omega (Sievers et al., 2011). Amino acid residues are labeled with asterisks (identical), colons (conserved) and dots (semiconserved). Secondary structure elements for the Cor a 1.04 isoforms are indicated on top. Hydrophobic, positively charged (donor), and negatively charged (acceptor) amino acid residues are indicated in yellow, blue and red, respectively. (B) Cor a 1.0401 structure (6Y3H) with the amphiphilic internal cavity colored as in (A), generated using Schrödinger’s Maestro Software Suite (Schrödinger, 2020). The positions of the six variable amino acid residues in the four Cor a 1.04 isoforms (4, 40, 62, 99, 130 and 158) and the cavity entrance ε1 are indicated. (C) Percentage of hydrophobic, proton donor, and proton acceptor interaction potential on the surface of the cavities in the four Cor a 1.04 isoforms.

One particular and conserved feature of PR-10 proteins is the large internal cavity formed by the secondary structure elements. The four Cor a 1.04 isoforms display cavity volumes of up to 1540 Å, which is within the range of other PR-10 allergens (Fernandes et al., 2013). In Cor a 1.0401 the surface of the internal cavity (Figure 2B) is formed by the hydrophobic residues Phe22 (α1), Ile30 (α2), Phe38 (L3), Ile56, Phe58, Ala 59 (β3), Phe64, Tyr66, Met67 (β4), Tyr83, Ile85 (β5), Gly88, Gly89, Pro90, Gly92 (L7), Ile98, Tyr100 (β6), Tyr121 (β7), and Ala136, Gly137, Leu144 (α3). An extended hydrophobic surface patch is located at the inner end of the cavity, where the three α-helices meet. In addition, a number of charged amino acid residues convey an amphiphilic character to the cavity, including residues His37 (L3), Glu60 (L5), Lys65, His69 (β4), Glu87 (β5), His93, Thr94 (L7), Asn130 (L9), and Glu132, Glu133, Lys140 (α3). This amphiphilic area extends to an entrance of the cavity (ε1), which is located between helix α3 and strands β3 and β4, as well as loops L5 and L6. The amphiphilic nature of the cavity has also been reported for other food and pollen allergens (Ahammer, Grutsch, Kamenik, Liedl, & Tollinger, 2017; Kofler et al., 2012; Orozco-Navarrete et al., 2020). Of note, the exact composition of the internal cavity in the four Cor a 1.04 isoforms is fairly diverse, in particular regarding the contributions from hydrophobic and charged residues (Figure 2 C). The percentage of hydrophobic surface area varies between 13 % and 20 %, while the contribution of positively charged residues to the cavity surface ranges from 32 % to 45 %, indicating a remarkable degree of variability between these proteins.

Plant food allergens from the PR-10 family typically bind natural flavonoids and other plant derivatives in the internal cavity (Fernandes et al., 2013). While natural ligands of the four hazelnut Cor a 1 allergens are not known to date, a flavonoid compound bound to hazel pollen PR-10 allergens was described recently. Structural data for hazel pollen PR-10 allergens are not available to date, but sequence comparison of nut and pollen allergens (63 % sequence identity, Figure 2A) suggests major differences regarding the internal surface composition. These include the exchange of charged and hydrophobic residues (H37A, G89D, T94K, Y100H, K140M) and charge inversion (H69E, H93D) of residues in the cavity. Nevertheless, the specific flavonoid bound to hazel pollen PR-10 allergens was shown to also bind to the hazelnut isoform Cor a 1.0401 *in vitro* (Jacob et al., 2019), implying that potential ligands bind to hazelnut and hazel pollen PR-10 proteins with relatively low specificity, in agreement with the current literature (Fernandes et al., 2013; Mogensen, Wimmer, Larsen, Spangfort, & Otzen, 2002). Moreover, the diverse nature of the internal cavities of the hazelnut Cor a 1.04 isoforms described above indicates binding of diverse ligands.

### The four Cor a 1.04 structures have different flexibilities

The NMR solution structural ensembles of the four Cor a 1.04 isoforms (Figure 1) suggest that only the C-termini and some loop regions of the four proteins are conformationally heterogeneous, in particular loop L5 connecting strands β3 and β4, which forms part of entrance ε1 to the internal cavity. A similar observation has previously been reported for other PR-10 food allergens, e.g. Mal d 1 (Ahammer et al., 2017), and it has been suggested that this segment of PR-10 proteins might function as a flexible gate keeper to the protein’s interior (Neudecker et al., 2001). For a more in-depth analysis of conformational heterogeneity in the four hazelnut Cor a 1.04 allergens we performed relaxation dispersion (RD) NMR experiments, which provide site-specific information about the presence of different conformers (Mittermaier & Kay, 2006). In RD-NMR experiments, transitions between different conformers occurring on the micro-to-millisecond time scale cause non-flat dispersion profiles, while conformational homogeneity results in flat dispersions. Experimental RD data for representative amino acid residues (Val23, Ser40/Gly40, Tyr100, and Thr118) in the four Cor a 1.04 isoforms are shown in Figure S2, and the corresponding RD amplitudes (ΔR_2,eff_ values) are color-coded on the protein structures in Figure 3. It is evident from the relaxation dispersion data that a significant portion of the protein backbone is flexible. Non-flat relaxation dispersion profiles are found for numerous residues in all four isoforms not only in loop regions, but also in all secondary structure elements. The most flexible regions include the short helices α1 and α2, parts of the β-sheet (β1, β5, and β7), and the middle of the C-terminal helix α3.

**Figure 3.**
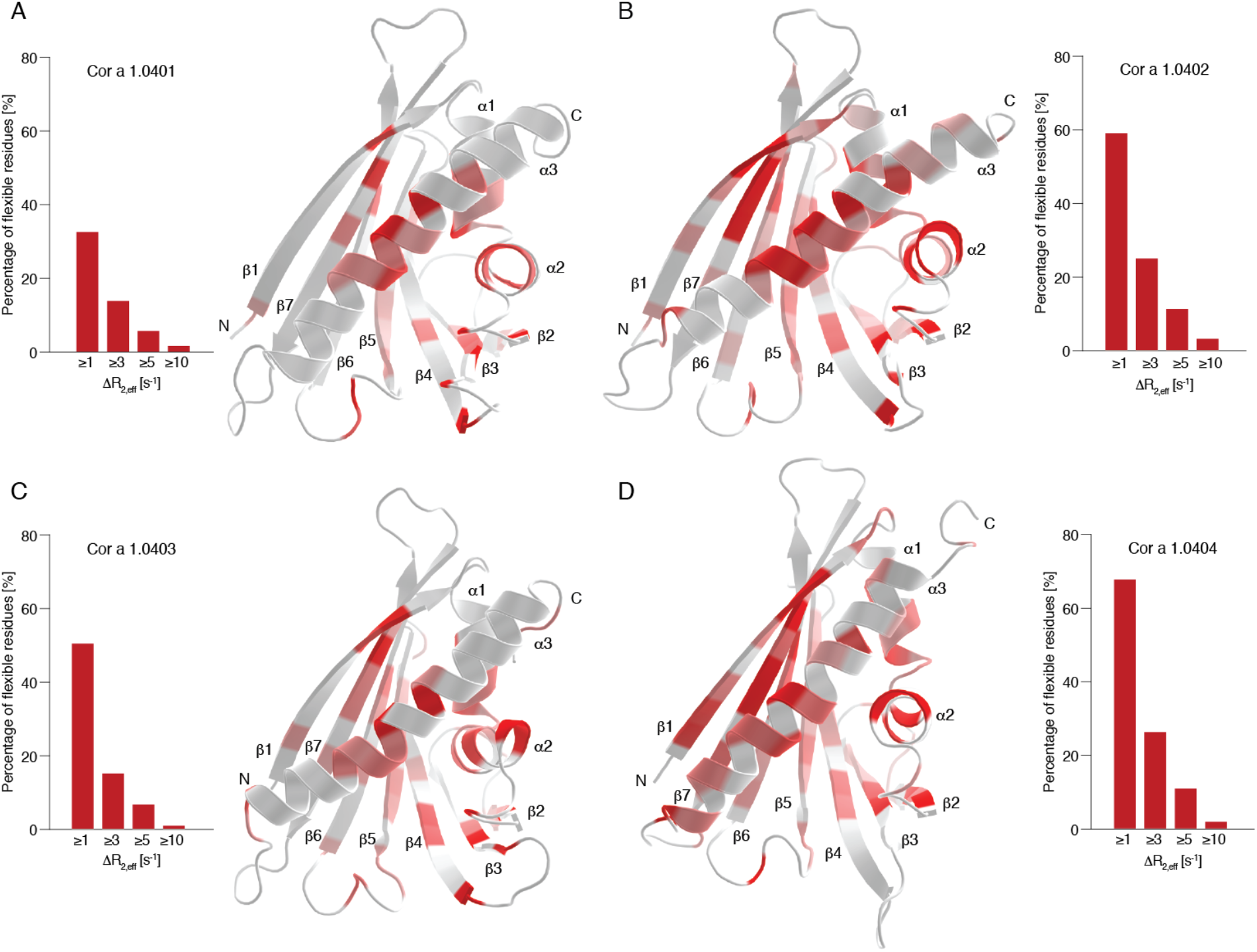
Structural flexibility of the four hazelnut isoforms Cor a 1.0401 (A), Cor a 1.0402 (B), Cor a 1.0403 (C), and Cor a 1.0404 (D). The bar plots show the percentage of flexible residues with relaxation dispersion amplitudes (ΔR_2,eff_ values) exceeding 1, 3, 5, and 10 s^-1^ at 600 MHz. ΔR_2,eff_ values plotted on the protein backbone with a color gradient from highly flexible (red) to rigid (white), using identical thresholds for all four proteins. Secondary structure elements and the N- and C-termini are indicated.

Interestingly, significant differences between the four isoforms are evident. The amplitudes of the RD-profiles exceed 1 s^-1^ for 33 %, 59 %, 50 %, and 68 % of all residues in Cor a 1.0401, Cor a 1.0402, Cor a 1.0403, and Cor a 1.0404, respectively (Figure 3). Likewise, RD amplitudes exceed 3 s^-1^ in 14 %, 25 %, 15 % and 26 % of all residues in these proteins, suggesting the “flexibility ranking” Cor a 1.0404 > Cor a 1.0402 > Cor a 1.0403 > Cor a 1.0401. Detailed comparison of the data clearly shows that the different flexibilities among the isoforms are not limited to the direct vicinity of the six variable amino acid residues. Indeed, for Cor a 1.0402 and especially for Cor a 1.0404 additional flexibility (when compared to the least flexible isoform Cor a 1.0401) is distributed almost over the entire protein scaffold. Moreover, it is evident that flexible residues are particularly grouped around position 99 in strand β6, which is occupied by proline in Cor a 1.0404 and serine in all other isoforms. This includes residues in the direct proximity of Pro99 in strand β6, such as Ile98 and Tyr100, as well as residues in the adjacent strands β5 and β7. Increased flexibility in β7 of Cor a 1.0404 is evident from both relaxation dispersion (Thr118 and Ser119) and NMR order parameter data (Ile114, Leu115, and Lys120, see Figure S3).

In isoforms Cor a 1.0401-03, the backbone amide of Ser99 forms a hydrogen bond to Lys120 in β7. It is likely that the lack of such a hydrogen bond in Cor a 1.0404 causes structural lability of the β-sheet of this particular isoform. Nevertheless, it is clear from the NMR relaxation data that even parts of the protein, which are distal from position 99 (*e*.*g*., helix α3), display the highest degree of flexibility in the isoform Cor a 1.0404.

### Cor a 1.0404 is structurally labile

The most (Cor a 1.0404) and the least (Cor a 1.0401) structurally flexible isoforms, have been reported as those with the lowest and the highest IgE-binding potential, respectively (Lüttkopf et al., 2002). Structural flexibility of proteins can result in transient exposure of backbone amides to solvent water, rendering them susceptible to exchange with surrounding water molecules. To further investigate structural flexibility in these two Cor a 1.04 isoforms, the solvent exposure of backbone amides was probed by use of NMR hydrogen-deuterium (H/D) exchange measurements that detect the replacement of the backbone amide protons with deuterium upon addition of D_2_O.

Figure 4 shows experimental hydrogen-deuterium exchange data for four exemplary amino acid residues Ala9, Glu101, Ala107, and Ala154. Because H/D exchange of most backbone amide protons in Cor a 1.0404 is remarkably fast, the experimental data for both isoforms were recorded at low temperature (10 °C) using rapid NMR data collection (SOFAST) techniques. ^1^H-^15^N correlation spectra were obtained in ca. 4 minutes, which enabled us to record reliable H/D exchange data, even for rapidly exchanging backbone amides, revealing significantly accelerated exchange in Cor a 1.0404 compared to Cor a 1.0401. In detail, Ala9, Glu101 and Ala154, which are located in secondary structure elements and hydrogen bonded, have very slow exchange rates in Cor a 1.0401, while Ala 107, which is surface exposed in loop L8, displays a moderate exchange rate. In Cor a 1.0404, all four amino acid residues display drastically faster H/D exchange. The positions of these residues are indicated on the Cor a 1.0404 structure in Figure 4, along with all other amino acid residues whose H/D exchange rate is at least four times faster in Cor a 1.0404 than in Cor a 1.0401 under identical conditions. Acceleration is not limited to the immediate sequential neighborhood of the two variable amino acid residues in the two isoforms (positions 4 and 99). Indeed, a number of residues that show increased exchange rates in Cor a 1.0404 are located in strand β7, which is adjacent to strand β6 containing Pro99.

**Figure 4.**
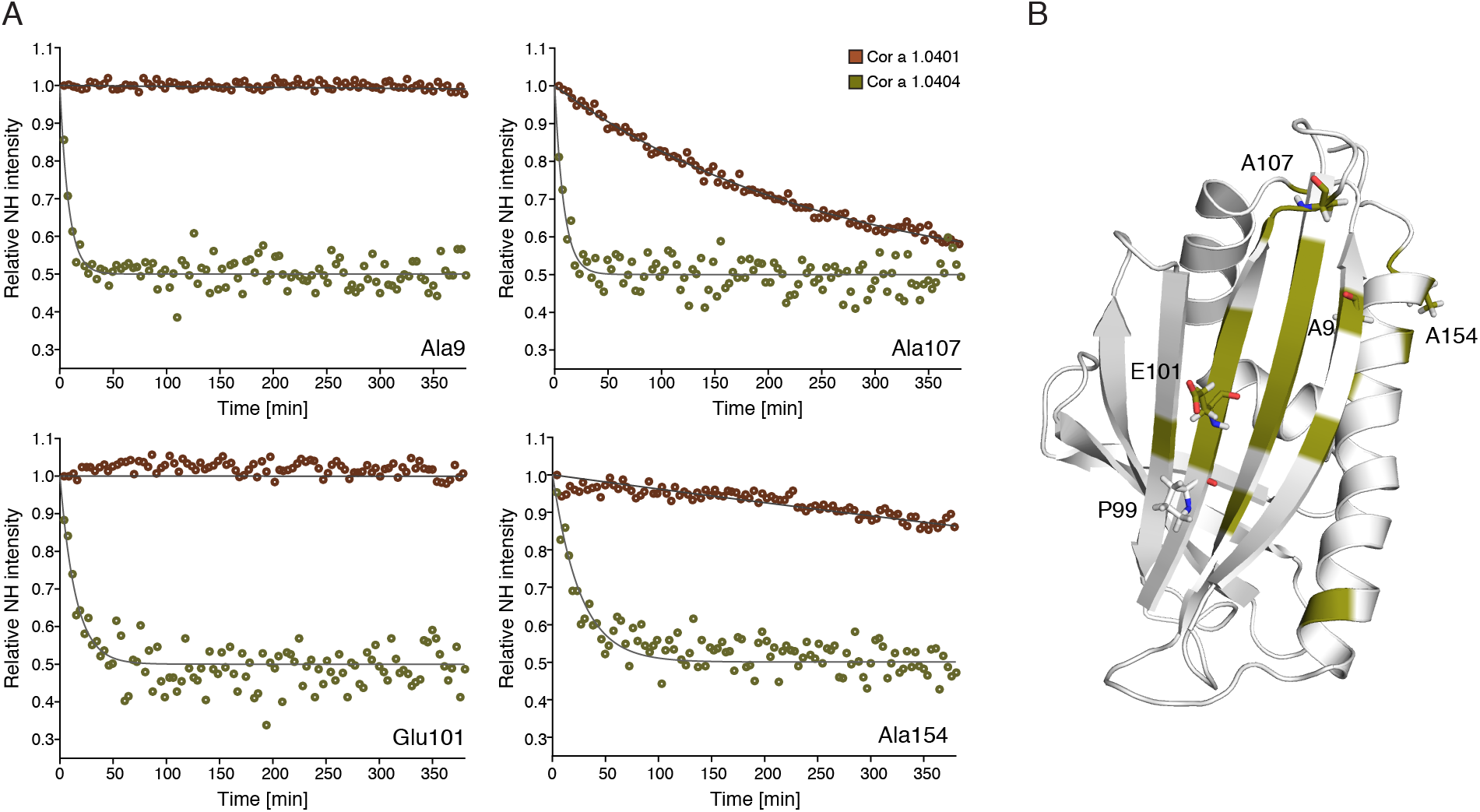
Hydrogen-deuterium exchange in Cor a 1.04 isoforms. (A) Time-dependent SOFAST ^1^H-^15^N-HMQC backbone amide intensities (circles), recorded at 10 °C, are shown for representative residues with drastically faster H/D exchange in Cor a 1.0404 (deep olive) than in Cor a 1.0401 (brown), along with best-fit exponential curves. (B) Amino acid residues for which H/D exchange is at least four times faster are highlighted in deep olive on the structure of Cor a 1.0404 (6Y3L). The four exemplary amino acid residues (Ala9, Glu101, Ala107, and Ala154) and Pro99 are displayed as sticks.

The fact that hydrogen-deuterium exchange of the two proteins Cor a 1.0401 and Cor a 1.0404, despite sharing similar structural scaffolds, exhibit significantly different exchange rates, indicates that hydrogen bonds in the Cor a 1.0404 isoform are weaker. This structural lability could be elicited by the missing hydrogen bond between Pro99 and Lys120 in the center of the β-sheet. Of note, the only other difference between the two isoforms is position 4 at the N-terminus, which is occupied by a cysteine in Cor a 1.0401 and a serine in Cor a 1.0404.

To investigate the impact of the two variable amino acid residues at positions 4 and 99 in Cor a 1.0401 and Cor a 1.0404 in detail, we performed time-dependent NMR degradation assays (Figure 5). After five and seven days at room temperature, Cor a 1.0404 shows degradation and aggregation peaks, while Cor a 1.0401 spectra remain unchanged during this time period. NMR resonance assignments of a partly degraded sample of Cor a 1.0404, identified the presence of small unstructured peptides Tyr5-Ile13 and Pro31-Ala34, indicating proteolytic cleavage of this particular isoform. Three mutant forms, C4S Cor a 1.0401, P99A Cor a 1.0404, and P99T Cor a 1.0404 were also investigated regarding degradation. While Cys4Ser probes the natural variability at position 4, the two proline mutants were chosen to examine the influence of Pro99. The NMR spectra of these variants show that their overall fold is not affected by the mutation and all three proteins were stable during the course of the assay (Figure S4). This implies that Pro99 in Cor a 1.0404 is the reason for the loss of stability.

**Figure 5.**
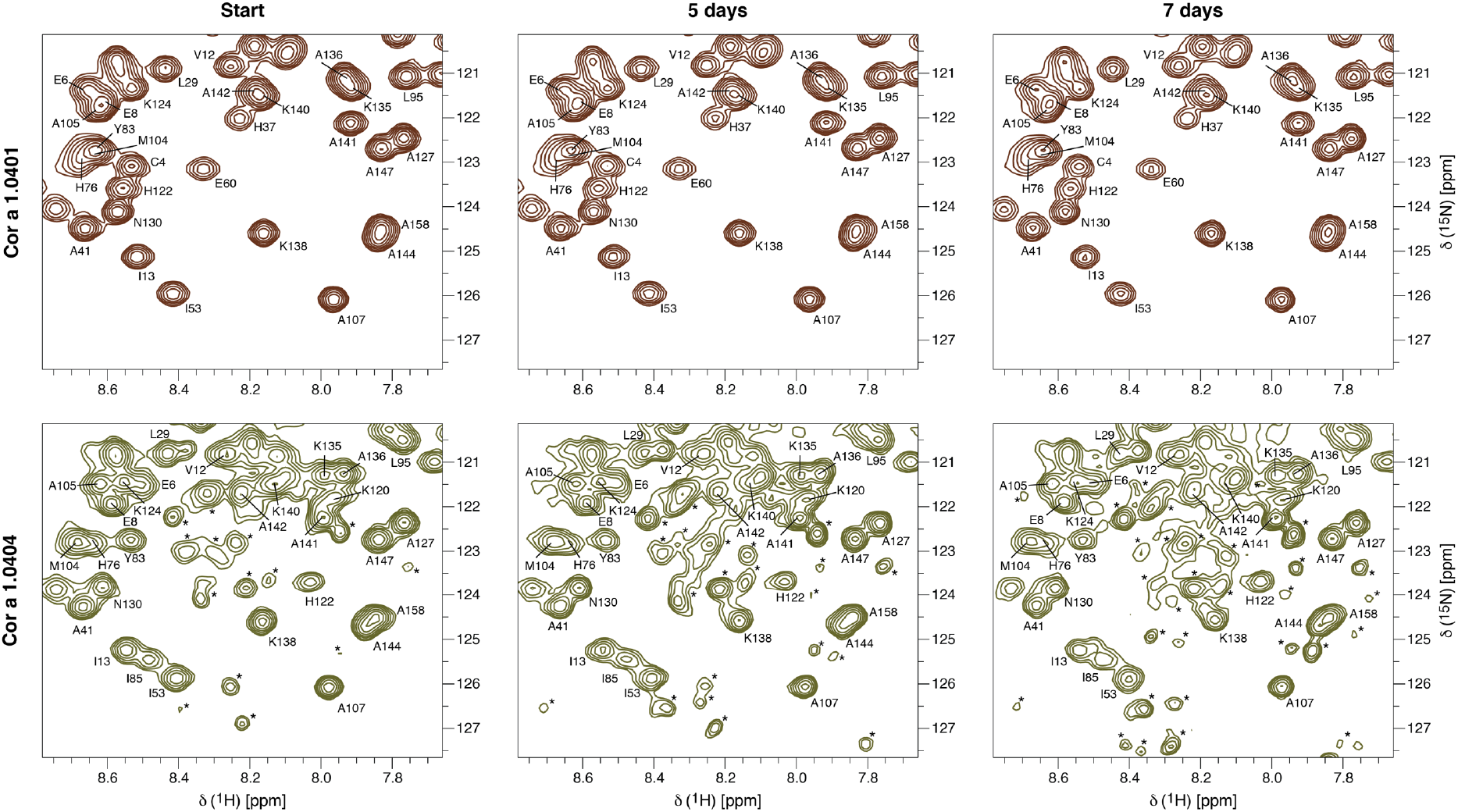
Time-dependent stability of Cor a 1.04 isoforms. Sections of ^1^H-^15^N-HSQC spectra of Cor a 1.0401 (top) and Cor a 1.0404 (bottom) directly after purification, as well as five and seven days later. Assignments (Führer et al., 2020) are indicated by single letter codes and signals labeled with an asterisk indicate aggregation and degradation.

Finally, the effect of temperature on Cor a 1.0401 was probed by acquiring ^1^H-^15^N-HSQC spectra at different temperatures in the range of 10 °C to 70 °C (Figure S5A). This protein remains folded even at 70 °C and precipitates at 80 °C. Heating of Cor a 1.0401 is fully reversible until 70 °C without intensity loss after cooling the sample again to 25 °C. This indicates high thermal stability of the Cor a 1.0401 structure in solution, in agreement with the observed IgE reactivity in roasted hazelnuts (Wigotzki, Steinhart, & Paschke, 2000). In contrast, for the isoform Cor a 1.0404 ^1^H-^15^N-HSQC spectra display substantially reduced intensities at temperatures above 40 °C, most likely due to accelerated chemical exchange of backbone amides with solvent (Figure S5B).

Taken together, our NMR data indicate general structural lability of the Cor a 1.0404 isoform that is very likely caused by the presence of Pro99 in the center of the β-sheet and a weakened hydrogen bonding network.

### Cor a 1.04 isoforms show different IgE-binding potentials

A previous study demonstrated binding of specific IgE from patients sera (Switzerland and Denmark) to the four hazelnut Cor a 1.04 isoforms (Lüttkopf et al., 2002). Measurably different IgE-binding properties were reported, with Cor a 1.0404 being the isoform with the lowest IgE-binding potential, while Cor a 1.0401 displayed the highest potential.

We verified the immunologic activity of our recombinantly produced Cor a 1.04 proteins using blood sera from twenty-two Austrian patients (Tyrol) included in a pilot study (Nothegger et al., 2020). The patients show birch-pollen related hazelnut allergy (specific IgE to Cor a 1 > 0.35 kU/mL and Bet v 1 > 0.35 KU/mL), positive skin-prick test to hazelnut extract and oral allergy symptoms to hazelnuts (see demographic data in Table 2). ELISA experiments reveal that Cor a 1.0401 indeed has the highest IgE-binding potential, while Cor a 1.0404 has the lowest. The isoforms Cor a 1.0402 and Cor a 1.0403 have moderate to high potentials, with the latter featuring slightly higher values, as evident from Figure 6A. In 91 % of all patients Cor a 1.0401 shows the highest binding of specific IgE, while Cor a 1.0404 barely shows binding potential. Besides, Cor a 1.0403 has a higher IgE-binding potential than Cor a 1.0402 in 91 % of all patients. Sera IgE reactivities toward the first three isoforms appear to be patient-specific, with considerable variations between them, while IgE reactivities toward Cor a 1.0404 are consistently low for all sera. Box plots of the patient-derived data confirm that specific IgE values for isoforms Cor a 1.0401-03 display relatively broad distributions within the patient group, whereas Cor a 1.0404 values are systematically low (Figure 6B). A statistical analysis of the ELISA data and ImmunoCAP data are shown in Figure 6C, indicating that the IgE-binding potential of Cor a 1.0401 shows no significant difference with the ImmunoCAP data and Cor a 1.0403. Additionally, no difference is observed for Cor a 1.0402 with Cor a 1.0403 and Cor a 1.0404. In summary, the patient derived ELISA data clearly suggest the immunologic ranking Cor a 1.0401 > 03 > 02 > 04.

**Table 2.** Demographic and immunologic characteristics of 22 patients with birch pollen-related hazelnut allergy.

| Characteristics | Values |
| --- | --- |
| Sex, n (f/m) | 22 (16/6) |
| **Median (range)** |  |
| Age [y] | 36 (22 - 68) |
| SPT [mm]* | 5.5 (3.0 - 9.0) n=13 |
| **ImmunoCAP** |  |
| IgE total [kU/L] | 124.0 (9.4 - 948.0) |
| Cor a 1.04 specific IgE [kU/L] | 8.02 (0.87 - 52.10) |
| Bet v 1 specific IgE [kU/L] | 14.40 (2.96 - 91.40) |
| **ELISA** |  |
| Cor a 1.0401 specific IgE [kU/L] | 2.02 (1.31 - 15.15) |
| Cor a 1.0402 specific IgE [kU/L] | 1.21 (0.82 - 4.98) |
| Cor a 1.0403 specific IgE [kU/L] | 1.57 (0.95 - 13.08) |
| Cor a 1.0404 specific IgE [kU/L] | 0.88 (0.48 - 1.41) |

\* Skin-Prick test (SPT) was determined as positive if the wheal was > 3 mm.

**Figure 6.**
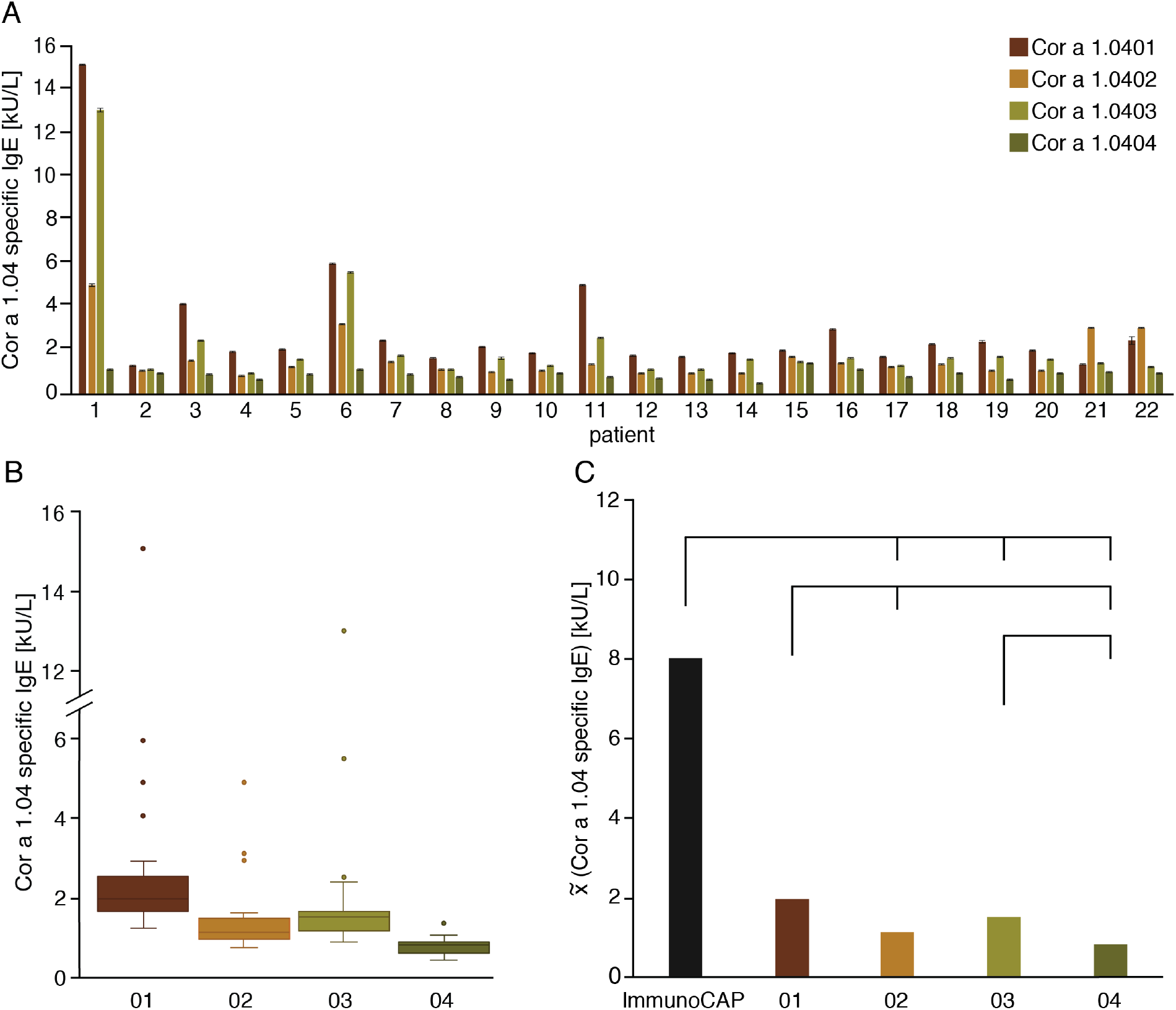
Specific IgE-binding to the four hazelnut isoforms Cor a 1.0401 (brown), Cor a 1.0402 (orange), Cor a 1.0403 (olive), and Cor a 1.0404 (deep olive). (A) Bar plots and standard deviations of IgE-binding potentials obtained by ELISA for each patient. (B) Box plots of the IgE-binding potentials of the four isoforms. The boxes are restricted by the 25^th^ and 75^th^ percentile, median values are given by a horizontal line inside the box and whiskers cover the minimal and maximal values inside the interquartile range (1.5 IQR). Patients with high total IgE levels and therefore high specific IgE are indicated by dots. (C) ELISA data (median values) and ImmunoCAP data (black). Differences between the IgE-binding potentials are given by the lines above with a p-value < 0.05 as statistically significant. The exact pairwise p-values are 0.006 (01-02), 0.453 (01-03), 1.89**·**10^−8^ (01-04), 0.565 (01-ImmunoCAP), 1.000 (02-03), 0.100 (02-04), 9.33**·**10^−7^ (02-ImmunoCAP), 0.001 (03-04), 0.001 (03-ImmunoCAP), and 2.44**·**10^−14^ (04-ImmunoCAP). All three panels suggest the immunologic ranking Cor a 1.0401 > 03 > 02 > 04.

As mentioned above, between Cor a 1.0401 and Cor a 1.0404, only two amino acid residues are different (Cys4/Ser4 and Ser99/Pro99), with their side-chains located on the protein surface, and our mutational studies suggest that proline at position 99 of Cor a 1.0404 causes the structural lability of this particular isoform. Likewise, Pro99 is presumably the main reason for the dramatically reduced IgE-binding potential of Cor a 1.0404 compared to Cor a 1.0401, as suggested by (Lüttkopf et al., 2002). This is corroborated by our IgE-binding data for the isoforms Cor a 1.0402 and Cor a 1.0403, which differ only at position 4 (Ser4/Cys4) and display comparable IgE-binding, with Cor a 1.0403 having only a slightly higher binding potential than Cor a 1.0402. These data suggest that replacement of a serine by a cysteine residue at position 4 has a minor effect on IgE-binding, even though these residues are surface exposed, as previously noted in enzyme allergosorbent tests (Lüttkopf et al., 2002).

In principle, IgE-binding can also be modulated by the formation of dimers or higher oligomers (Kofler et al., 2014). Cor a 1.04 isoforms contain between two and three cysteine residues (at positions 4, 82 and 160) that are surface exposed and accessible for cysteine-mediated dimerization. Therefore, we probed the oligomerization state of Cor a 1.0401 (three cysteines) and Cor a 1.0404 (Cys82 and Cys160) under the buffer conditions that we employed for our ELISA and NMR experiments using pulsed-field gradient experiments. Cor a 1.0401 has a hydrodynamic radius of 18.6 ± 0.1 Å without and 18.7 ± 0.4 Å with a reducing agent (DTT) being present, respectively, which is consistent with a monomeric protein in both cases and comparable to other monomeric PR-10 proteins (Ahammer et al., 2017; Grutsch et al., 2014). This is supported by the retention times observed in size exclusion chromatography, which are characteristic for a monomeric PR-10 protein. Likewise, the hydrodynamic radius of Cor a 1.0404 is 18.6 ± 0.5 Å, again indicating a monomeric protein. Cor a 1.04 isoforms thus show no tendency for dimerization or oligomerization under the experimental conditions that were used for IgE-binding measurements.

## Conclusions

Our NMR experimental and immunologic data suggest an inverse relationship between the IgE-binding potential and the structural flexibility of the four known isoforms of the Cor a 1 hazelnut allergens, even though their three-dimensional structures are highly similar. Cor a 1.0401 is the isoform with the highest potential to bind IgE, while it is the protein in this group with the most rigid backbone scaffold. Cor a 1.0404, on the other hand, is the isoform that has the lowest IgE-binding potential, yet this protein is strikingly flexible and conformationally heterogeneous in solution and displays a weak hydrogen bonding network. Cor a 1.0402 and Cor a 1.0403 are intermediate between these isoforms regarding their IgE-binding potential and their structural flexibility.

Enhanced levels of structural flexibility have been observed for various allergens (Neudecker et al., 2004), including those from the PR-10 family of proteins (Grutsch et al., 2017; Moraes et al., 2018). Only for the major birch pollen allergen, Bet v 1, isoform-specific flexibility data have been reported, revealing Bet v 1.0101 to be fairly rigid and having substantially higher IgE-binding potential than the structurally flexible isoform Bet v 1.0102 (Grutsch et al., 2017; Wagner et al., 2008). As with Cor a 1, structural flexibility in Bet v 1 is distributed across the entire PR-10 scaffold, including secondary structure elements and loops. These observations parallel our experimental data for the hazelnut allergen described here.

Structural flexibility has been recognized as a critical component of antigen-antibody binding (Fernandez-Quintero et al., 2019). Indeed, from structural studies it is evident that antibody affinity maturation is, in part, driven by the rigidification of the antigen-binding site (Mishra & Mariuzza, 2018). The reduction of antibody structural flexibility was proposed as means to lower the entropic cost of complex formation in order to design antibodies with increased affinities (Yanaka, Moriwaki, Tsumoto, & Sugase, 2017). Along these lines, it has been shown by experiment that complex formation leads to antibody rigidification (Yanaka et al., 2017), and reduced antibody flexibility has been proposed as general feature of antigen-antibody binding (Williams, Rule, Poljak, & Benjamin, 1997)

Experimental data regarding the rigidification of allergens in antibody complexes are not available to date. Considering the pronounced flexibility of PR-10 allergens in solution, it is possible that complex formation is indeed accompanied by rigidification of the protein scaffold, resulting in a loss in conformational entropy and causing a lower binding affinity for more flexible isoforms. According to the concept of conformational selection, it is also conceivable that in the more rigid isoforms the binding-competent conformation is highly populated in solution. The more flexible isoforms on the other hand may fluctuate between several conformational states, resulting in a lower population of the binding competent conformation. This lack of structural pre-organization could contribute to lower IgE-binding. The inverse relation between structural flexibility and IgE-binding observed in our Cor a 1 study suggests that entropic contributions indeed play an appreciable role in complex formation between these allergens and antibodies.

## Materials and Methods

### Plasmid generation and protein expression

The construction of plasmids encoding the hazelnut allergens Cor a 1.0401 (accession no. **AAD48405**), Cor a 1.0402 (accession no. **AAG40329**), Cor a 1.0403 (accession no. **AAG40330**), and Cor a 1.0404 (accession no. **AAG40331**) and the subsequent expression has been described previously (Führer et al., 2020). Plasmid constructs encoding the mutant forms C4S Cor a 1.0401, P99A Cor a 1.0404, and P99T Cor a 1.0404 were created by site-directed mutagenesis using Phusion DNA polymerase (New England Biolabs, Frankfurt am Main, Germany) for C4S Cor a 1.0401 and Platinum SuperFi DNA polymerase (Thermo Fisher Scientific, Vienna, Austria) for P99A Cor a 1.0404, and P99T Cor a 1.0404 according to the protocol of the manufacturers. The mutant forms P99A Cor a 1.0404 and P99T Cor a 1.0404 were generated using C4S Cor a 1.0401 as template. The used oligonucleotides were 5’-ccatgggcgtgttc<u>tcc</u>tacgaagatgagg-3’ and 5’-cctcatcttcgta<u>gga</u>gaacacgcccatgg-3’ for C4S Cor a 1.0401, 5’-ccctggaaaaaatc<u>gcc</u>tacgagattaaaatggc-3’ and 5’-gccattttaatctcgta<u>ggc</u>gattttttccaggg-3’ for P99A Cor a 1.0404, and 5’-ccctggaaaaaatc<u>acc</u>tacgagattaaaatggc-3’ and 5’-gccattttaatctcgta<u>ggt</u>gattttttccaggg-3’ for P99T Cor a 1.0404 (underlined nucleotides were mutated). Mutagenesis was verified by DNA-sequencing. Expression of the three mutant proteins was performed as for the wild-type hazelnut isoforms but without using ISOGRO^®^-^15^N powder for the expression (Führer et al., 2020). Unlabeled proteins were expressed in M9 minimal medium without isotopically labeled reagents.

### Protein purification and preparation

The protein purification protocol for the four wild-type hazelnut isoforms has been described previously (Führer et al., 2020). The same protocol was used for the three mutant forms. All NMR samples contained 20 mM sodium phosphate buffer (pH 6.9), 2 mM DTT, 9 % D_2_O and 0.5 mM ^15^N labeled or ^15^N/^13^C labeled protein.

### Mass spectrometry experiments

For the determination of the accurate mass of the hazelnut Cor a 1.04 isoforms a 7 Tesla Fourier-transform ion cyclotron resonance (FT-ICR) mass spectrometer (Apex Ultra 70, Bruker Daltonics, Bremen, Germany), equipped with an electrospray ionization (ESI) source was used. Unlabeled Cor a 1.0401, Cor a 1.0402, Cor a 1.0403, and Cor a 1.0404 protein samples were desalted five times with ca. 500 µL 100 mM ammonium acetate pH 6.8 and subsequently five times with ca. 500 µL H_2_O. Afterwards, the samples were diluted with H_2_O/CH_3_OH (1:1, v/v) supplemented with 1 % acetic acid to ca. 1 µM. Data analysis was done with Bruker Compass DataAnalysis and Bruker Daltonics Bio Tools or mMass (Strohalm, Kavan, Novak, Volny, & Havlicek, 2010) (version 3.0.0).

### NMR structure determination

The resonance assignments for the four isoforms with all corresponding NMR experiments have been reported previously and are available at the Biological Magnetic Resonance Data Bank (https://bmrb.wisc.edu) under the accession numbers 27965, 27961, 27967, and 28016 for Cor a 1.0401, Cor a 1.0402, Cor a 1.0403, and Cor a 1.0404, respectively (Führer et al., 2020). For structure determination, three-dimensional ^1^H-^15^N-NOESY-HSQC, ^1^H-^13^C-NOESY-HSQC, and aromatic ^1^H-^13^C-NOESY-HSQC experiments with a mixing time of 150 ms each were carried out at 25 °C on a 500 MHz Agilent DirectDrive 2 spectrometer (Agilent Technologies, Santa Clara, CA, USA) equipped with a room temperature probe. Processing of the NMR data was performed with NMRPipe (Delaglio et al., 1995), and spectra were visualized with nmrDraw (Delaglio et al., 1995) and analyzed using CcpNmr (Vranken et al., 2005).

For the assignment of the NOE cross-peaks a structure model generated by the program CS-Rosetta (Shen et al., 2010; Shen et al., 2008; Shen, Vernon, Baker, & Bax, 2009) was used and the obtained cross-peaks were converted into distance restraints, based on their intensities. These restraints were further categorized by their intensities ranging from very strong to very weak with upper limits of 2.8 Å, 3.0 Å, 5.0 Å, 6.0 Å, and 6.5 Å. Hydrogen bond restraints were derived from the CS-Rosetta structure model for the secondary structure elements, if no water cross-peak was detected in the proton TOCSY spectra. Dihedral angle restrains (Φ and Ψ) were derived from the program TALOS+ (Shen, Delaglio, Cornilescu, & Bax, 2009).

For the structure determination, the program XPLOR-NIH (version 2.52) (Schwieters, Kuszewski, & Clore, 2006; Schwieters, Kuszewski, Tjandra, & Clore, 2003) was used. Initially, 500 structures were calculated in 3000 steps at an initial temperature of 7000 K, followed by 10000 cooling steps. The 20 structures with the lowest energy were further refined in explicit solvent. For this purpose, each structure was surrounded by a cubic box of TIP3P (Jorgensen, Chandrasekhar, & Madura, 1983) waters with at least 10 Å from each atom to the box boundaries. The LEaP module of AMBER 18 (D.A. Case, 2018) using the Amber force field 14SB (Maier et al., 2015) parameterized the system. After solvent relaxation (Wallnoefer, Liedl, & Fox, 2011), simulated annealing calculations with a Langevin thermostat (Adelman & Doll, 1976) (collision frequency: 2 ps^-1^) and a Berenden barostat (Berendsen, Postma, van Gunsteren, DiNola, & Haak, 1984) (relaxation time: 2 ps) were performed. Hydrogen bonds were constrained with the SHAKE algorithm (Ciccotti & Ryckaert, 1986) and a van der Waals cutoff of 10 Å was used along with the particle-mesh Ewald method (Darden, York, & Pedersen, 1993) for long range electrostatics. A simulated annealing scheme of 50 ns with a time step of 1 fs for each structure was performed using the NOE distance restraints. The protein structure validation software (PSVS) suite (Bhattacharya, Tejero, & Montelione, 2007) was used to validate the refined structures. Internal cavity volumes and surface hydrophobicity were determined using SiteMap (T. Halgren, 2007; T. A. Halgren, 2009) as implemented in the Schrödinger Maestro Software Suite (Schrödinger, 2020). RMSD values between the different PR-10 proteins were obtained using the program SuperPose (Maiti, Van Domselaar, Zhang, & Wishart, 2004).

### Temperature dependency and NMR diffusion experiments

The temperature sensitivity of Cor a 1.0401 and Cor a 1.0404 was probed by recording ^1^H-^15^N-HSQC spectra on a 700 MHz Bruker Avance Neo spectrometer (Bruker BioSpin, Rheinstetten, Germany) equipped with a Prodigy CryoProbe at different temperatures (10, 15, 20, 25, 30, 35, 40, 50, 60, 70 °C). Chemical shifts were referenced at all temperatures using an external standard sample containing 1 % (w/v) 4,4-dimethyl-4-silapentane-1-sulfonic acid (DSS) in 450 µL 20 mM sodium phosphate buffer (pH 6.9), 2 mM DTT, supplemented with 10 % D_2_O.

For the determination of the oligomerization state of Cor a 1.0401 and Cor a 1.0404 stimulated echo pulsed field gradient experiments were used (Choy et al., 2002). The employed gradient field strengths were 2.0, 4.5, 7.0, 9.5, 12.0, 14.5, 17.0, 19.5. 22.0, and 24.5 G/cm with a constant diffusion time of 160 ms. Hydrodynamic radii were determined for all dispersed peaks in the ^1^H-^15^N-HSQC spectrum by using an in-house MATLAB fitting program and the Stokes-Einstein equation, as described in detail for Bet v 1 (Grutsch et al., 2014). Average values and standard deviations of the hydrodynamic radii were calculated from the 20 residues with the lowest RMSD values.

### NMR hydrogen-deuterium exchange experiments

Hydrogen-deuterium exchange of the amide protons of Cor a 1.0401 and Cor a 1.0404 was assessed by SOFAST ^1^H-^15^N-HMQC spectra (Schanda, Forge, & Brutscher, 2007) acquired at 700 MHz using a Prodigy CryoProbe at 10 °C and a rapid injection device, as described (Schanda, Brutscher, Konrat, & Tollinger, 2008). NMR samples were prepared in 20 mM sodium phosphate buffer (pH 6.9) with 2 mM DTT, and hydrogen-deuterium exchange was initiated by addition of an equal volume of the same buffer in D_2_O to a final protein concentration of 0.3 mM. A series of SOFAST ^1^H-^15^N-HMQC spectra were recorded immediately after sample mixing, each lasting for 4.12 minutes (Cor a 1.0401) or 3.8 minutes (Cor a 1.0404). Using an in-house MATLAB fitting script, the hydrogen-deuterium exchange rates were obtained from exponential decay fits.

### NMR relaxation experiments

Backbone amide ^15^N relaxation dispersion experiments were recorded on a 500 MHz spectrometer, a 600 MHz Bruker Avance II+ spectrometer (Bruker BioSpin, Rheinstetten, Germany) equipped with a Prodigy CryoProbe, and a 700 MHz spectrometer, using sensitivity enhanced Carr-Purcell-Meiboom-Gill (CPMG) sequences (Loria, Rance, & Palmer, 1991; Tollinger, Skrynnikov, Mulder, Forman-Kay, & Kay, 2001) with ^1^H continuous-wave decoupling during the CPMG period (D. F. Hansen, Vallurupalli, & Kay, 2008). Spectra were recorded at different CPMG field strengths ν_CPMG_=(2τ_CPMG_)^-1^, with τ_CPMG_ being the time between two consecutive 180° pulses in the CPMG pulse train. The ν_CPMG_ field strengths were 33.3, 66.7, 100.0, 133.3, 166.7, 200.0, 266.7, 333.3, 466.7, 600.0, 733.3, and 933.3 Hz, with repeat experiments at 66.7 Hz and 600.0 Hz, and the length of the CPMG pulse train was set to T_relax_ = 30 ms in all experiments. Effective relaxation rates R_2,eff_ = -1/T_relax_ · ln(I/I_0_) were determined from partial peak volumes (intensities in 5 x 5 grids) of the resonances in the individual spectra (I), along with a reference intensity (I_0_) with T_relax_ set to 0. The so-obtained relaxation dispersion profiles were analyzed by fitting the exact CPMG expression for two-site exchange using an in-house-written MATLAB script (Baldwin, 2014). Per-residue relaxation dispersion amplitudes (ΔR_2,eff_) were calculated from the 600 MHz data as ΔR_2,eff_ = R_2,eff_ (ν_CPMG_ = 0) - R_2,eff_ (ν_CPMG_ = ∞), where R_2,eff_ (ν_CPMG_ = ∞) and R_2,eff_ (ν_CPMG_ = 0) are the extrapolated R_2,eff_ values at infinite and zero CPMG field strengths, respectively.

Experiments for measuring longitudinal and rotating-frame relaxation rates (R_1_ and R_1ρ_) and the ^15^N{^1^H} steady-state NOE (nuclear Overhauser effect) (Akke & Palmer, 1996; Barbato, Ikura, Kay, Pastor, & Bax, 1992; Farrow et al., 1994; Korzhnev, Orekhov, & Kay, 2005) for Cor a 1.0401 and Cor a 1.0404 were performed at 500 MHz. Relaxation periods for measuring R_1_ rates were 11.1, 55.5, 111.0, 222.0, 333.0, 444.0, 555.0, and 666.0 ms with repeat experiments at 55.5 ms and 333.0 ms. For measuring R_1ρ_ rates, relaxation periods were 10.0, 20.0, 30.0, 40.0, 50.0, 60.0, 70.0, and 90.0 ms, with repeats at 20.0 ms and 50.0 ms. For ^15^N{^1^H} steady-state NOE measurements two spectra were recorded, one with an interscan delay (d1) of 3 s and proton saturation for 1.25 s and the second experiment with d1 set to 4.25 s without proton saturation. The relaxation rates R_1_ and R_1ρ_ were determined from exponential fits using an in-house MATLAB fitting script. Transverse relaxation rates R_2_ were derived from R_1_ and R_1ρ_ values as described (Trott & Palmer, 2002) and order parameters S^2^ were determined with the program FAST-Modelfree (version 4.15) (Cole & Loria, 2003), which has the model-free approach implemented (Clore et al., 1990; Lipari & Szabo, 1982a, 1982b).

### IgE-binding experiments

A skin-prick test (SPT) with hazelnut extract (ALK-Abelló, Linz, Austria), histamine as a positive control and diluent as a negative control was performed on the flexor surface of the forearm according to Heinzerling et al. (L. Heinzerling et al., 2013; L. M. Heinzerling et al., 2009). Serum levels of total IgE and IgE specific for Bet v 1 and Cor a 1 were determined by ImmunoCAP (Phadia®250, Thermo Fisher Scientific, Uppsala, Sweden) according to the manufacturer’s specifications. The demographical and immunologic data (Table 2) are expressed as median values with corresponding ranges.

The IgE-binding potential and the biological activity of Cor a 1.0401, Cor a 1.0402, Cor a 1.0403, and Cor a 1.0404 was assessed by using an enzyme-linked immunosorbent assay (ELISA). Unlabeled freshly recombinantly produced proteins and the mAb 107 of the Human IgE ELISA development kit HRP (Mabtech, Nacka Strand, Sweden), which was used for standardization, were diluted in 0.1 M NaHCO3 pH 9.7 to a final concentration of 1 µg/mL. In each well of a 96 Well Corning Costar Assay Plate (Merck, Darmstadt, Germany) either 100 µL mAB 107 (for standardization) or 100 µL protein (for specific IgE detection) were placed and incubated at 4 °C overnight. The plate was washed with 150 µL PBS/0.1 % Tween 20 the next day and subsequently saturated with 150 µL PBS/3 % BSA per well. Incubation was conducted for 90 minutes under moderate shaking and afterwards the plate was washed as before. The patients’ blood sera were diluted 1:2 (v/v) with PBS and the human IgE standard from the kit was diluted to 100, 50, 20, 10, 5, 2, and 0 ng/mL with PBS. Either 50 µL of diluted serum or diluted standard were added to the wells containing protein or mAb 107, respectively and the plate was incubated at 4 °C overnight. Another washing step was conducted as before and the Anti-Human IgE (ε-chain specific) detecting antibody (Merck, Darmstadt, Germany) was diluted 1:1000 (v/v) with PBS/1 % BSA and 50 µL of it were added to each well. Incubation was conducted for 60 minutes and afterwards the plate was washed as before and 100 µL SureBlue TMB Microwell Peroxidase Substrate (SeraCare, Milford, MA, USA) were added to each well and incubated for 20 minutes in the dark. The reaction was stopped by the addition of 100 µL TMB Stop Solution (SeraCare, Milford, MA, USA). Between each incubation step the plate was sealed with adhesive plate sealers (Biozol, Eching, Germany) to avoid evaporation and contamination. The absorption at 650 nm was measured at a microplate reader (BMG Labtech, Mitterdorf a. d. Raab, Austria). Each protein and standard was measured as triplicate and a linear logarithmic function was derived from the standards. Values for Cor a 1.04 specific IgE [kU/L] was determined using this linear function.

The Shapiro-Wilk test was used to evaluate normal distribution. Correlations between specific Cor a 1.04 IgE values with the IgE values determined by ImmunoCAP were assessed by the Spearman rank test. Differences between the groups were tested with the Friedman two-way ANOVA and Post hoc tests. All statistical analyses were done with the program SPSS 25 (IBMCorp., 2017).

## Acknowledgements

We thank Dr. Katrin Breuker, Dr. Thomas Müller, Christina Meisenbichler, and Michael Palasser for mass spectrometry experiments and Jana Unterhauser for assistance with ELISA experiments. This work was supported by the Austrian Science Fund FWF (P26849 to MT, P30737 to KRL), and the Austrian Research Promotion Agency FFG (West Austrian BioNMR 858017). Patients’ blood sera were provided by the Department of Dermatology, Venerology and Allergology at the Medical University of Innsbruck within the AppleCare Study (funded by the European Regional Development Fund Interreg V-A Italy-Austria 2014-2020).

