## Supplementary Material for "Structural basis for variable IgE reactivities of Cor a 1 hazelnut allergens"

**Figure S1**

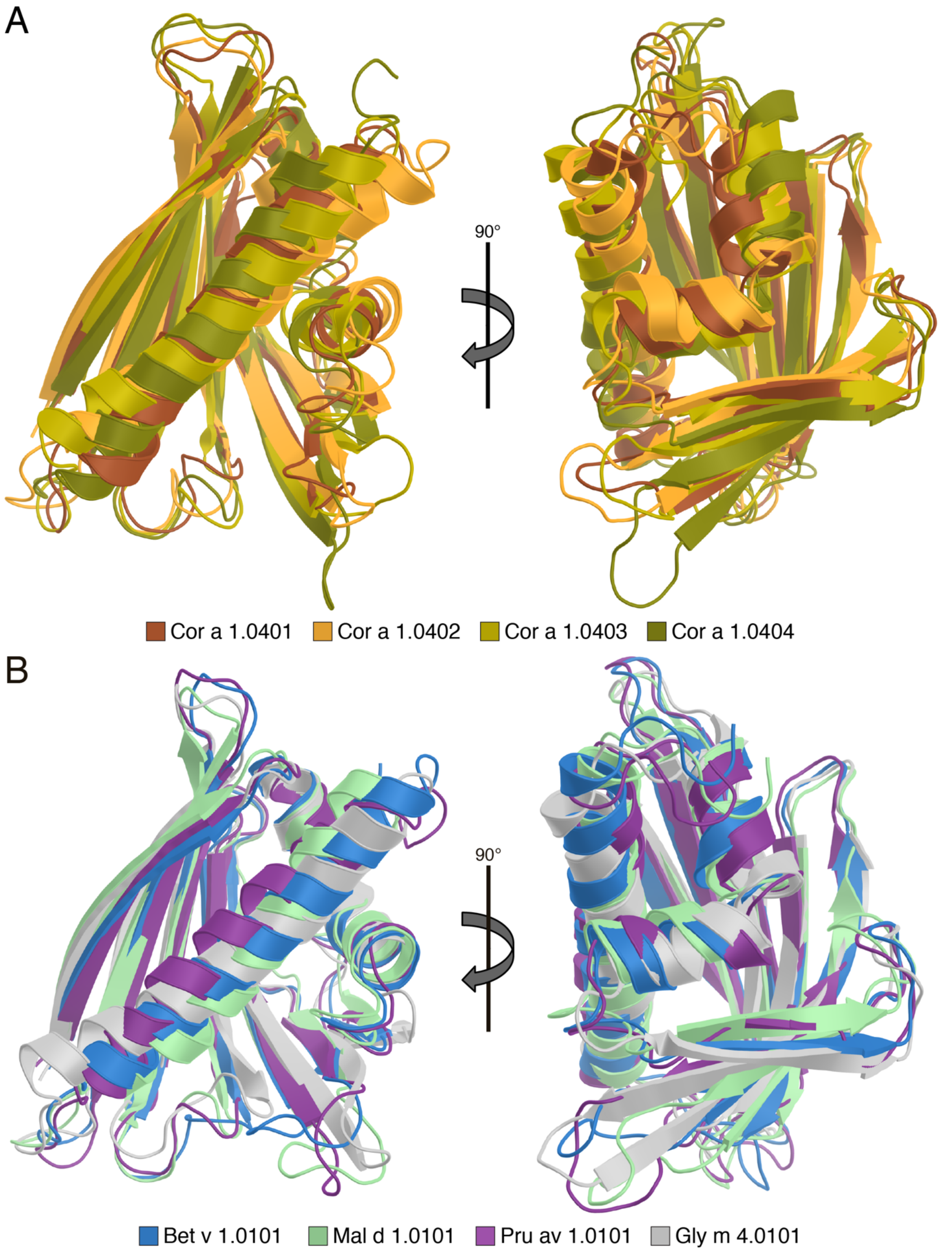


Structural comparison of the four hazelnut Cor a 1.04 isoforms with other PR-10 allergens. (A) Overlay of the solution structures of Cor a 1.0401 (brown, 6Y3H), Cor a 1.0402 (orange, 6Y3I), Cor a 1.0403 (olive, 6Y3K), and Cor a 1.0404 (deep olive, 6Y3L). (B) Overlay of the structures of Bet v 1.0101 (blue, 4A88), Mal d 1.0101 (pale green, 5MMU), Pru av 1.0101 (purple, 1E09), and Gly m 4.0101 (gray, 2K7H).

#### Figure S2

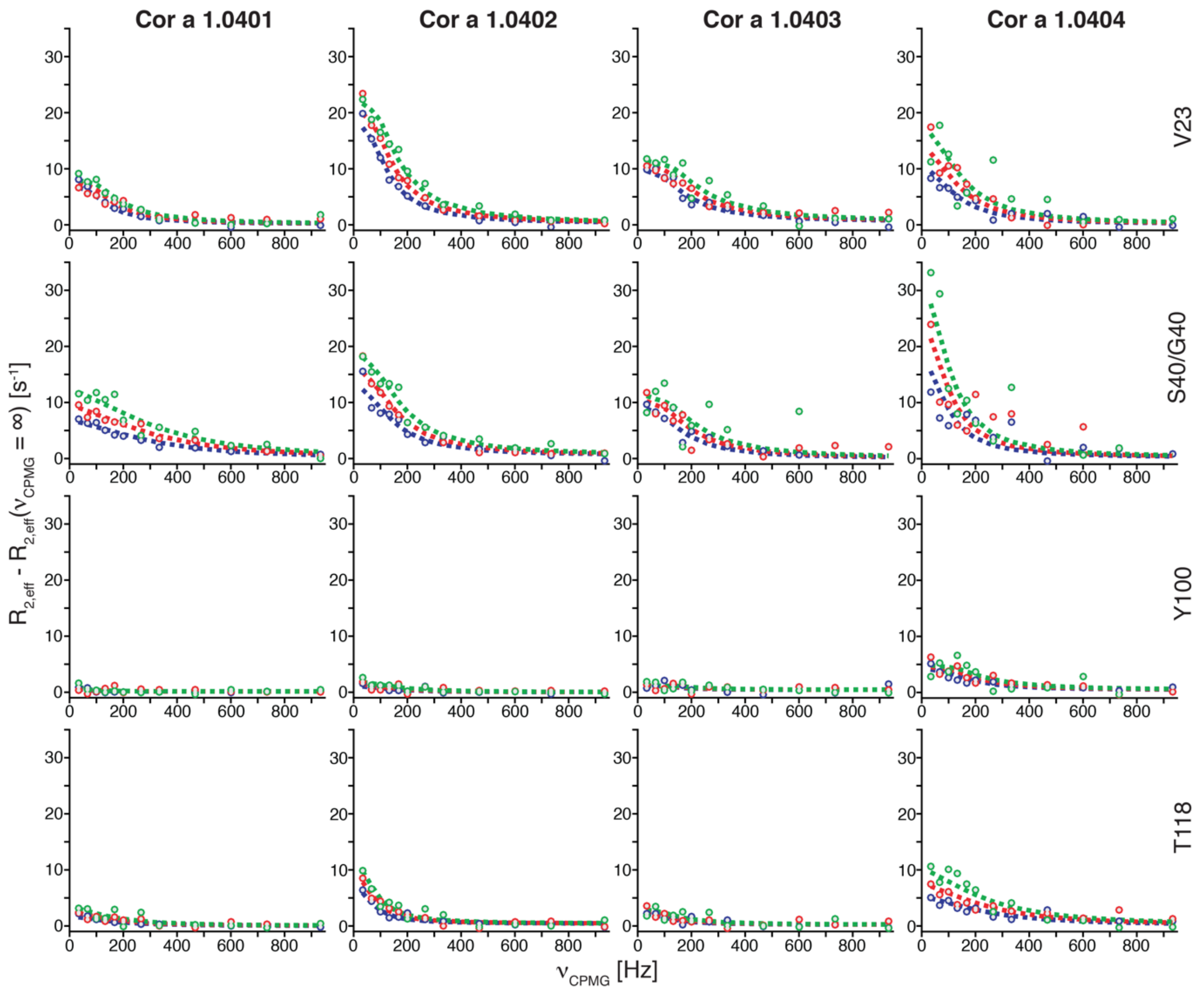


Backbone amide NH relaxation dispersion profiles of the four hazelnut Cor a 1.04 isoforms. $R_{2,eff}$ - $R_{2,eff}$ ($\nu_{CPMG} = \infty$) values at variable CPMG field strengths $\nu_{CPMG}$, acquired at 500 MHz (blue), 600 MHz (red), and 700 MHz (green) are shown along with best-fit dashed lines. Data are shown for four representative amino acid residues in the short helix α1 (Val23), in strand β2 (Ser40 in Cor a 1.0401 and Cor a 1.0404, Gly40 in Cor a 1.0402 and Cor a 1.0403) and in the vicinity of position 99 in strands β6 (Tyr100) and β7 (Thr118). Relaxation dispersion amplitudes ($\Delta R_{2,eff}$) used in Figure 3 are given by $R_{2,eff} - R_{2,eff}(\nu_{CPMG} = \infty)$ at $\nu_{CPMG} = 0$.

**Figure S3**

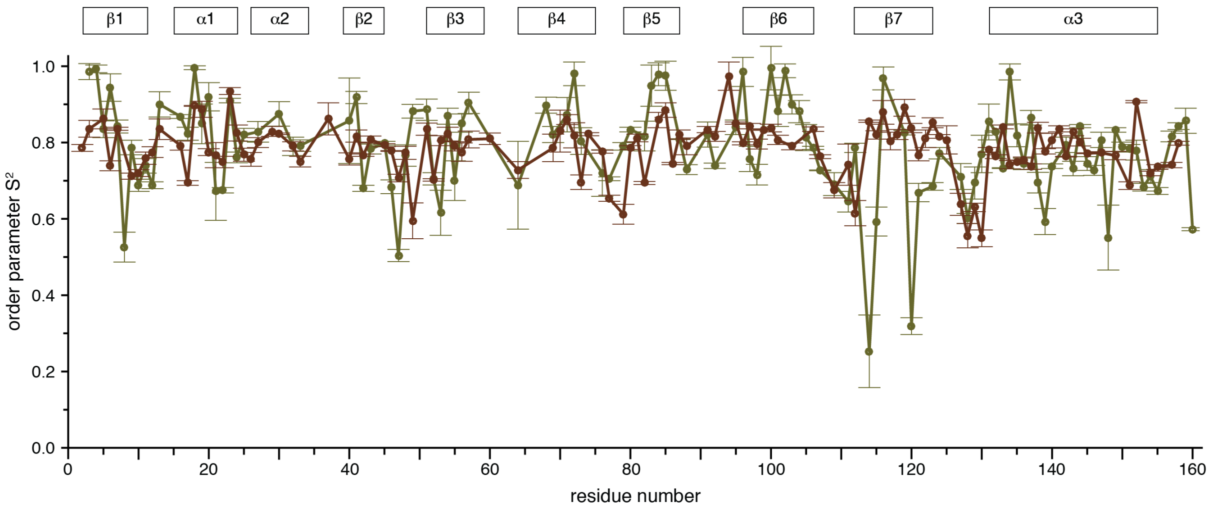


NMR-derived backbone amide NH order parameters $S^2$ of Cor a 1.0401 (brown) and Cor a 1.0404 (deep olive), obtained with the program FAST-Modelfree (Cole & Loria, 2003). Residue specific $S^2$ values are depicted as circles and error bars are shown. Solid lines connecting residues for which experimental data are available are drawn for better visualization. Low order parameters indicate high flexibility on the picosecond to nanosecond time scale. Secondary structure elements of the hazelnut Cor a 1.04 isoforms are shown on top.

### Figure S4

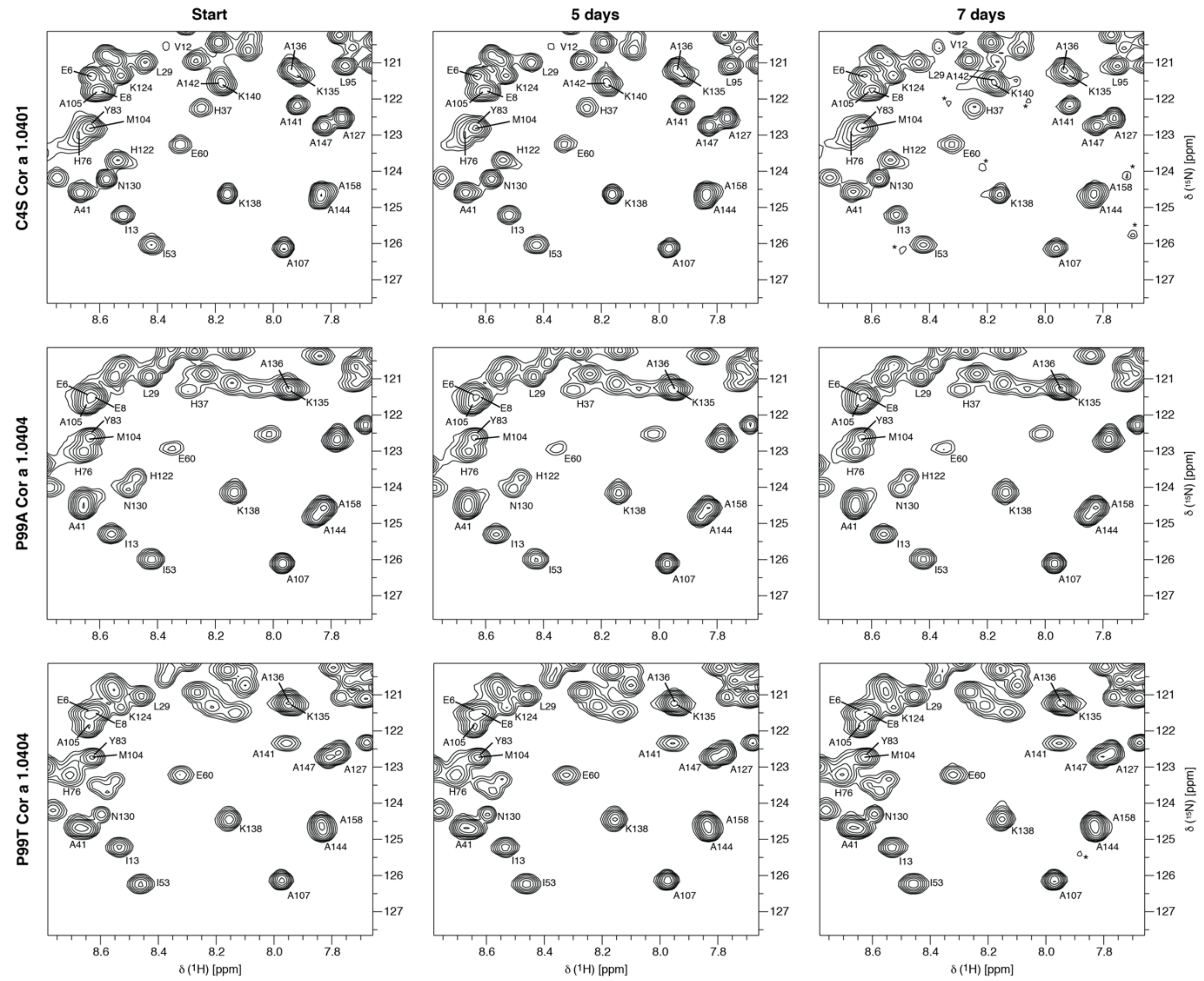


Time-dependent stability of three mutant forms of Cor a 1.0401 and Cor a 1.0404. Sections of $^{1}$H-$^{15}$N-HSQC spectra of C4S Cor a 1.0401 (top), P99A Cor a 1.0404 (middle), and P99T Cor a 1.0404 (bottom) directly after purification, as well as five and seven days later. Tentative assignments are indicated by single letter codes and signals labeled with an asterisk indicate aggregation and degradation.

**Figure S5**

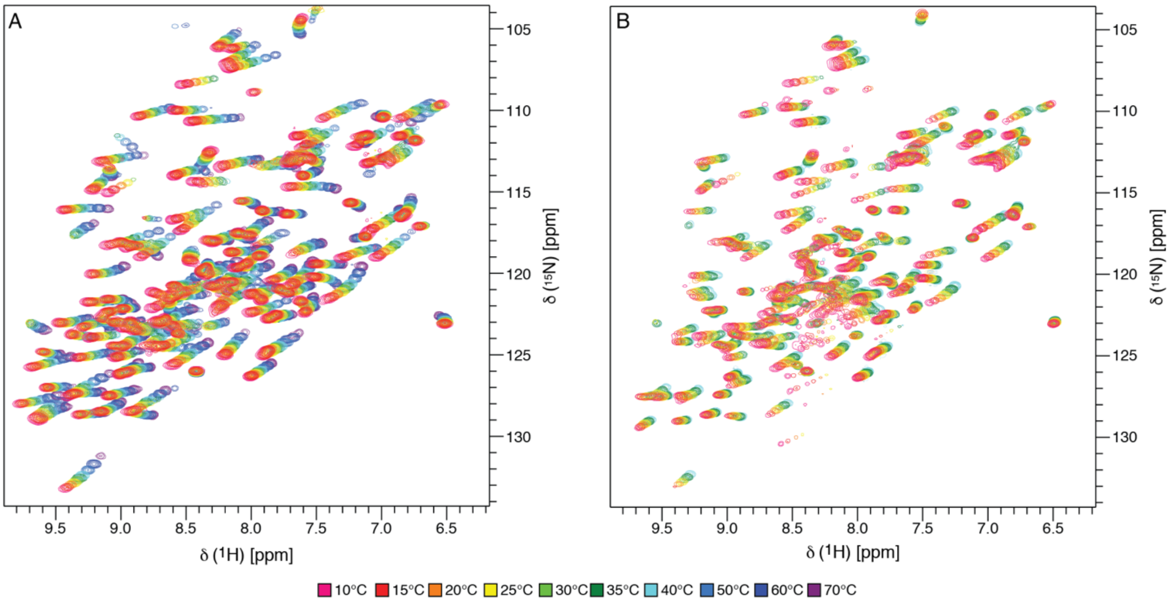


Temperature-dependent stability of Cor a 1.04 isoforms. Temperature dependent 700 MHz $^{1}H$-$^{15}N$-HSQC spectra of Cor a 1.0401 (A) and Cor a 1.0404 (B) are superimposed (referenced to DSS). Cor a 1.0401 spectra could be acquired up to 70 °C, while Cor a 1.0404 resonances vanished above 40 °C.
